# Preservation of HIV-1 Gag helical bundle symmetry by bevirimat is central to maturation inhibition

**DOI:** 10.1101/2021.05.24.445455

**Authors:** Alexander J. Pak, Michael D. Purdy, Mark Yeager, Gregory A. Voth

## Abstract

The assembly and maturation of human immunodeficiency virus type-1 (HIV-1) requires proteolytic cleavage of the Gag polyprotein. The rate-limiting step resides at the junction between the capsid protein CA and spacer peptide 1, which assembles as a six-helix bundle (6HB). bevirimat (BVM), the first-in-class maturation inhibitor drug, targets the 6HB and impedes proteolytic cleavage, yet the molecular mechanisms of its activity, and relatedly, the escape mechanisms of mutant viruses, remain unclear. Here, we employed extensive molecular dynamics (MD) simulations and free energy calculations to quantitatively investigate molecular structure-activity relationships, comparing wild-type and mutant viruses in the presence and absence of BVM and inositol hexakisphosphate (IP6), an assembly cofactor. Our analysis shows that the efficacy of BVM is directly correlated with preservation of six-fold symmetry in the 6HB, which exists as an ensemble of structural states. We identified two primary escape mechanisms, and both lead to loss of symmetry, thereby facilitating helix uncoiling to aid access of protease. Our findings also highlight specific interactions that can be targeted for improved inhibitor activity and support the use of MD simulations for future inhibitor design.

## Introduction

The late stages of human immunodeficiency virus type-1 (HIV-1) replication involve the formation of particles that subsequently undergo maturation to generate infectious virions (*1, 2*). Prior to maturation, group specific antigen (Gag) polyproteins are attached to the inner leaflet of the viral envelope in a pleomorphic, spherical, and quasi-hexagonal lattice (*3-5*). HIV-1 protease cleaves Gag between its domains during maturation, in which the rate-limiting cleavage resides between the capsid (CA) and spacer peptide 1 (SP1) (*6*). In the mature infectious particle, the released CA molecules assemble into a cone that encloses the viral nucleoprotein complex. The body of the cone is formed by CA hexamers that pack with quasi-hexagonal symmetry, and canonically, the wide and narrow ends of the cone are capped with seven and five CA pentamers, respectively (*7-10*).

Inhibition of HIV-1 maturation is the target of a novel class of small molecule drugs. bevirimat (BVM), or 3-O-(3’,3’-dimethylsuccinyl)betulinic acid, was the first proposed maturation inhibitor and acts by blocking cleavage at the helical junction between the C-terminal domain of CA (CA_CTD_) and SP1 (*11*). However, clinical trials were halted due to loss of susceptibility in over half of patients (*12, 13*). *In vitro* studies have shown that point mutations throughout the CA/SP1 junction confer BVM resistance, suggesting that the polymorphisms in this region, and in particular in the QVT motif of SP1, affect the mechanism of action of BVM (*14, 15*).

The current mechanistic model for BVM proposes that it stabilizes the CA/SP1 junction. Recent structural studies using X-ray crystallography (*16*), cryo-electron tomography (cryo-ET) (*17*), and electron diffraction of frozen-hydrated 3D microcrystals (MicroED) (*18*) resolved this junction as a six-helix bundle (6HB). Intriguingly, the six scissile bonds (between L363 and A364 in each protomer) face the interior of the bundle in which the central pore sterically occludes access of HIV-1 protease. Cryo-electron microscopy (cryo-EM) and magic angle spinning nuclear magnetic resonance analysis of CA/SP1 protein tubules, which adopt mature-like conformations, suggests that the CA/SP1 junction exists in a dynamic helix-coil equilibrium in the mature state (*19*). Cryo-ET analysis of a series of Gag cleavage mutants similarly showed mature lattices with disordered CA/SP1 junctions (*20*). However, immature lattices maintained ordered, helical CA/SP1 junctions (*20*). It is therefore unclear whether immature Gag lattices undergo a dynamic helix-coil equilibrium, and if this equilibrium is necessary for protease activity and relatedly, affected by BVM.

The recent high-resolution (2.9 Å) MicroED structure of BVM-bound CA_CTD_/SP1 revealed that a single BVM molecule stabilizes the 6HB via both electrostatic interactions with the dimethylsuccinyl moiety and hydrophobic interactions with the pentacyclic triterpenoid ring of BVM (*18*). Interestingly, a comparison between the drug-free and BVM-bound structures suggests that no major structural changes throughout the 6HB are induced by BVM. Likewise, recent solid-state nuclear magnetic resonance (ssNMR) experiments confirmed that BVM-induced structural changes are subtle, which further suggested that there are minimal changes in the conformation of the 6HBs (*21*). From these experiments, it can be inferred that the 6HBs may not exist in a dynamic helix-coil equilibrium. Consequently, the molecular mechanisms driving BVM activity and, relatedly, how escape mutants recover viral activity remain unclear. More recently, cryo-EM and MD analysis suggested that the edges of the fissures within immature HIV-1 capsid particles expose incomplete Gag hexamers, which could be the potential substrates for protease cleavage (*22*). This would imply that only the protomers within incomplete 6HBs may be in a dynamic helix-coil equilibrium.

Given our understanding of the chemical interactions between BVM and CA_CTD_/SP1 (*18*), we sought to further address the detailed mechanism of action of BVM by performing all-atom molecular dynamics (MD) simulations of BVM bound to wild type (WT) Gag CA_CTD_/SP1 and three CA_CTD_/SP1 mutations that impart varying degrees of drug resistance – L363, A364V, and V370A, the latter of which is part of the polymorphic QVT motif (*14, 15*). We find that BVM binds to the interior of the 6HB in two binding poses that productively rigidify the 6HB. The 6HB remains largely α-helical and rigidification occurs with respect to fluctuations about the six-fold symmetry axis of the bundle, which is cooperatively enhanced by the presence of inositol hexakisphosphate (IP6), an assembly cofactor (*23*). Resistance mutants escape BVM activity under two modes: abrogating knob-in-hole interactions between helices or promoting non-uniform BVM-6HB interactions. In both cases, the six-fold symmetry of the bundle is perturbed and facilitates helix-to-coil transitions, thereby enabling proteolytic cleavage and maturation, as demonstrated by our free energy calculations. Our findings highlight that preservation of symmetry throughout the ensemble of 6HB structural states is an important characteristic of maturation inhibitor activity.

## Results

### Characterization of the six-helix bundle conformational ensemble

We first analyzed the impact of BVM binding on the conformational ensemble of the 6HB. We prepared MD simulations of a solvated CA_CTD_/SP1 hexamer (PDB: 5L93 (*17*)) with restraints to helix 9 in order to emulate the quaternary structure of the immature lattice (see Methods). As seen in Figure 1, BVM was positioned in the center of the 6HB. Two binding poses were initialized – the dimethylsuccinyl moiety pointing toward the K359 ring (or “BVM-up”) and pointing toward the C terminus (or “BVM-down”). We simulated three configurations of WT virus (drug-free, BVM-up, and BVM-down), both with and without IP6, each with five independent MD replicas of 0.60 µs. We used the same procedure to simulate the escape mutants L363F, A364V, and V370A. The aggregate MD statistics (across 120 trajectories) encompassed 72 µs.

**Figure 1.**
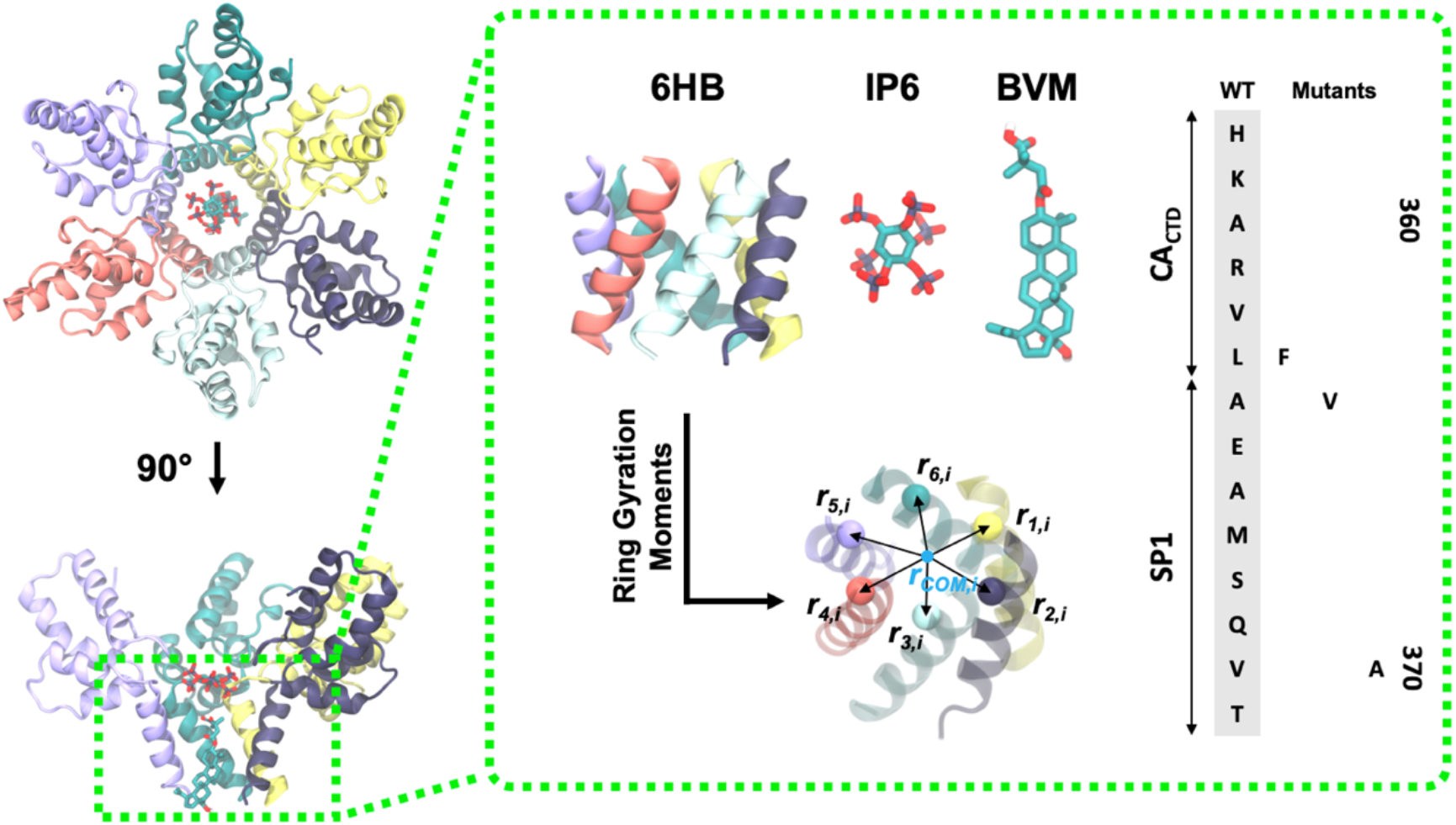
Schematic of HIV-1 CA_CTD_/SP1 with inositol hexakisphosphate (IP6) and bevirimat (BVM). The highlighted portion depicts the six-helix bundle (6HB), which contains the binding sites for IP6 and BVM; two of the protomers are omitted in the side view for clarity. The conformational ensemble of the 6HB was assessed using ring gyration moments (see main text) using the bond vectors shown in the schematic. The WT sequence spanning the CA_CTD_/SP1 junction is shown (H358 to T371), as well as the three mutants (L363F, A364V, and V370A) investigated in this work.

To characterize the 6HB conformation, we used a combination of dimensional reduction techniques. We described each 6HB configuration by a high-dimensional set of quantitative descriptors, a procedure known as featurization. First, we computed gyration moments on a per-ring basis along the 6HB (see Figure 1). For each residue with index *i* between H356 and T371, we calculated the mean (*µ*_*i*_) of the distance between each of the six Cα atoms of residue *i* (*r*_*k,i*_) and their collective center of mass (*r*_*com,i*_):

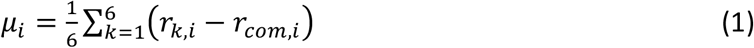

We then calculated the second through fourth moments about *µ*_*i*_, i.e. *µ*_*2,i*_ through *µ*_*4,i*_:

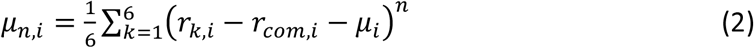

Therefore, for each 6HB configuration, a total of 64 features were collected. Our use of features that represent the collective configuration of the six helices ensures permutation invariance across helix labels. The features from all 120 trajectories were used for dimensional reduction.

Dimensional reduction was performed using time-lagged independent component analysis (tICA) (*24-26*), a linear projection technique that aims to maximize representation of auto-correlations (see Methods). The largest eigenmodes correspond to the slowest varying changes within the constraints of linear transforms of the feature set, and have been used to describe collective variables of biomolecules. Here, we find that the first two tICs capture 25% of the kinetic variance. It is therefore evident that the 6HB conformational space is not easily projected onto a simple 2D manifold in tIC space. Instead, we find that the first ten components (tICs) capture up to 84% of the kinetic variance. We then clustered the 10D tIC-projected data using k-means clustering (*27*) and filtered the data to 7 clusters with the highest populations, representing 76% of all configurations (see Methods). For visualization purposes, the aggregated data projected onto these ten tICs were embedded onto a 2D manifold using multidimensional scaling (*28*). Figure 2A depicts the filtered and labeled tIC data on the embedded 2D manifold.

**Figure 2.**
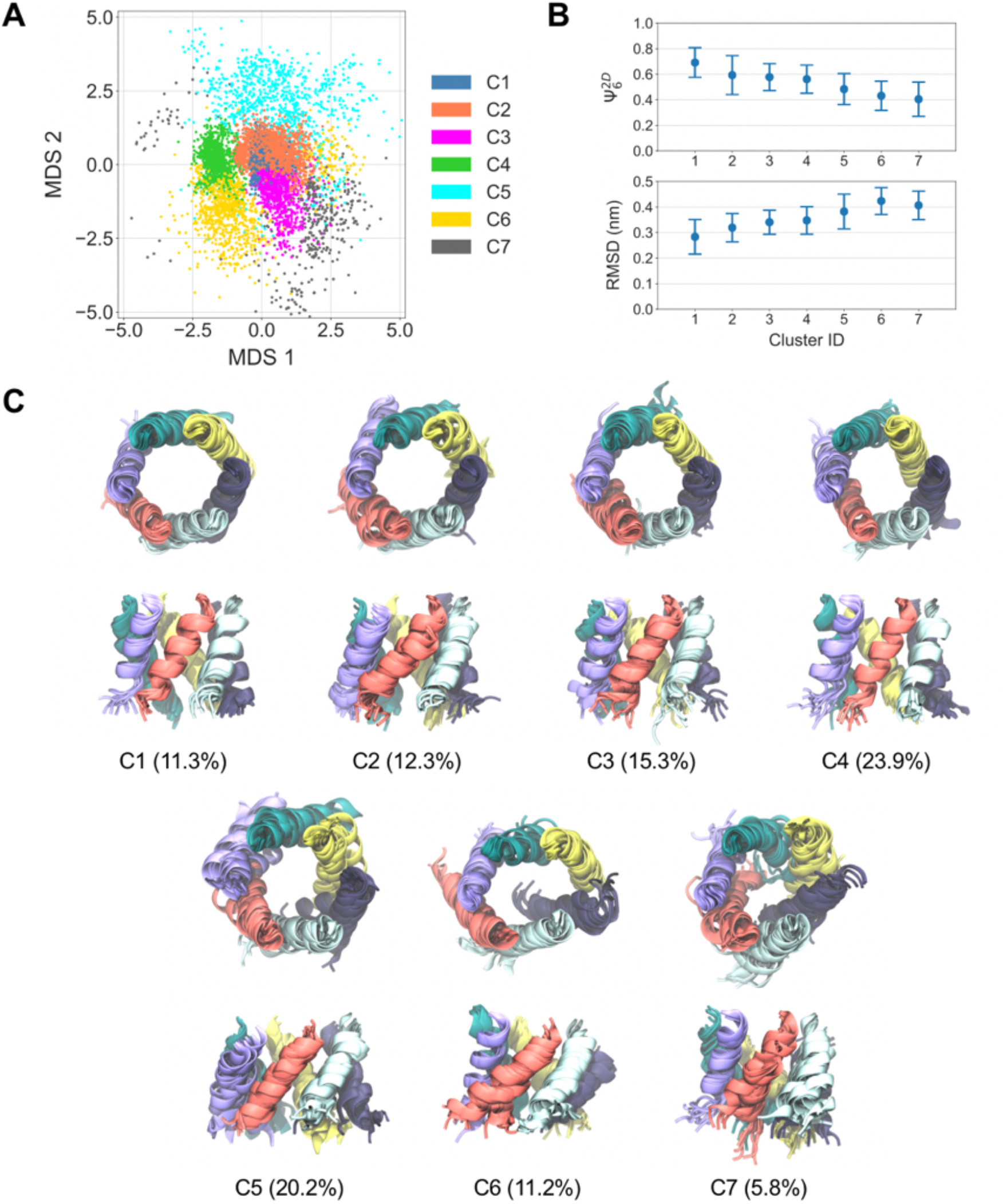
Clustering of six-helix bundle (6HB) configurational states. (A) Two-dimensional projection of the aggregate ten-dimensional time-lagged independent components using multidimensional scaling (MDS). Data points are colored according to their cluster index from k-means clustering. A randomly selected subset (n=7200; 2% of dataset) of the data is shown here for clarity. (B) Comparison of the 2D six-fold Mermin order parameter (Ψ_6_^2D^) and root-mean-squared-deviation (RMSD) of all 6HB configurations within each cluster; cluster labels were sorted in decreasing order of Ψ_6_^2D^ (n=360,000). (C) Representative snapshots of the 6HB structure classified by each cluster. Each cluster shows top and side views of 10 randomly selected and aligned configurations with each helix colored separately. The percentage of the total population of the data that is represented by each respective cluster is shown in parentheses.

Each of the 7 filtered clusters represents a distinct subpopulation of 6HB states, which were analyzed further. For each configuration within each cluster, we computed the root-mean-squared-deviation (RMSD) of the 6HB Cα backbones with respect to the atomic model (PDB 5L93 (*17*)). We also quantified the symmetry throughout the 6HB using the 2D six-fold Mermin order parameter(*29*) (Ψ_6_^2D^) for each C_α_ ring along the 6HB as follows:

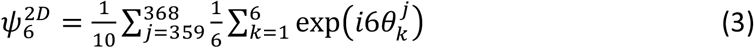

where θ^j^_k_ is the angle formed between the C_α_ atom of residue *j* in helix *k*, the center of mass (COM) of all C_α_ atoms of residue j, and the C_α_ atom of residue *j* in helix *k*+1. In other words, Ψ_6_^2D^ measures the average hexatic order on a per-ring basis throughout the 6HB; Ψ_6_^2D^ is expected to be a value of 1 if the 6HB is six-fold symmetric and diminishes toward 0 with increasing asymmetry. Finally, we quantified the similarity of the C_α_ backbone to an ideal α-helix (α_sim_) using the following:

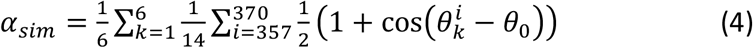

where θ^i^_k_ is the angle formed between the C_α_ of residue *i*-1, *i*, and *i*+1 in helix *k* and θ_0_ is the expected angle for α-helices (≅ 91°). When the 6HB is α-helical, α_sim_ is expected to be 1 and diminishes toward 0 with increasing dissimilarity.

Figure 2B compares the Ψ_6_^2D^ and RMSD across the 7 clusters, which were sorted with respect to the mean Ψ_6_^2D^. We find that decreasing Ψ_6_^2D^ across clusters is commensurate with increasing RMSD, i.e., the C1, C2, and C3 clusters correspond to the most symmetric conformations. However, α_sim_ remains around 0.96 in all clusters (see Supplementary Figure 1). These results suggest that the 6HB remains largely α-helical and fluctuations from the average 6HB structure, i.e., cluster 1, are largely due to distortions to the 6HB symmetry.

Representative configurations from each cluster are depicted in Figure 2C. Cluster 1 contains the most similar structure to the atomic model of the 6HB, which is both α-helical and six-fold symmetric. Clusters 2 to 4 are partially six-fold symmetric. Here, loss of six-fold symmetry is due to a combination of helical splaying (e.g. cluster 2) and contraction along lower-symmetry axes (e.g. cluster 4). Additional losses of six-fold symmetry are seen in clusters 5 to 7, which are ascribed to the disassociation of knob-in-hole interactions between adjacent helices. In the sections below, we analyze the role of BVM, IP6, and escape mutants on the expression of these clusters.

### bevirimat preserves while escape mutants disrupt symmetry in the six-helix bundle

Using our cluster definitions, we now quantify cluster populations in WT HIV-1 viruses, in which we compare both BVM-up and BVM-down binding orientations. As seen in Figure 3A, the cluster with the majority population (around 90%) in WT virus without BVM is cluster 3, i.e., 6HBs with partial six-fold symmetry; the minority population (around 10%) is cluster 6, i.e., asymmetric 6HBs. Interestingly, cluster 1, i.e., 6HBs with six-fold symmetry, is negligibly populated in drug-free WT virus. The presence of BVM, however, broadly shifts the cluster populations in favor of clusters 1 to 5. In the BVM-up orientation, the population of cluster 1 is around 40%, and more populous than that of the BVM-down orientation (around 25%). The primary difference between the two binding poses is the association between the dimethylsuccinyl moiety and the K359 ring in the BVM-up case, which was previously proposed on the basis of MicroED structural studies (*18*). This stabilization occurs even though the distance between the carboxylate of dimethylsuccinyl and the ammonium of K359 is greater than is typically seen for salt bridges, which may arise due to the stoichiometry of six lysines to one dimethylsuccinul in BVM (*18*). Our analysis suggests that this interaction is productive for six-fold symmetry preservation in the 6HB, but both poses appear to be viable binding modes and each restricts the loss of symmetry in the 6HB.

**Figure 3.**
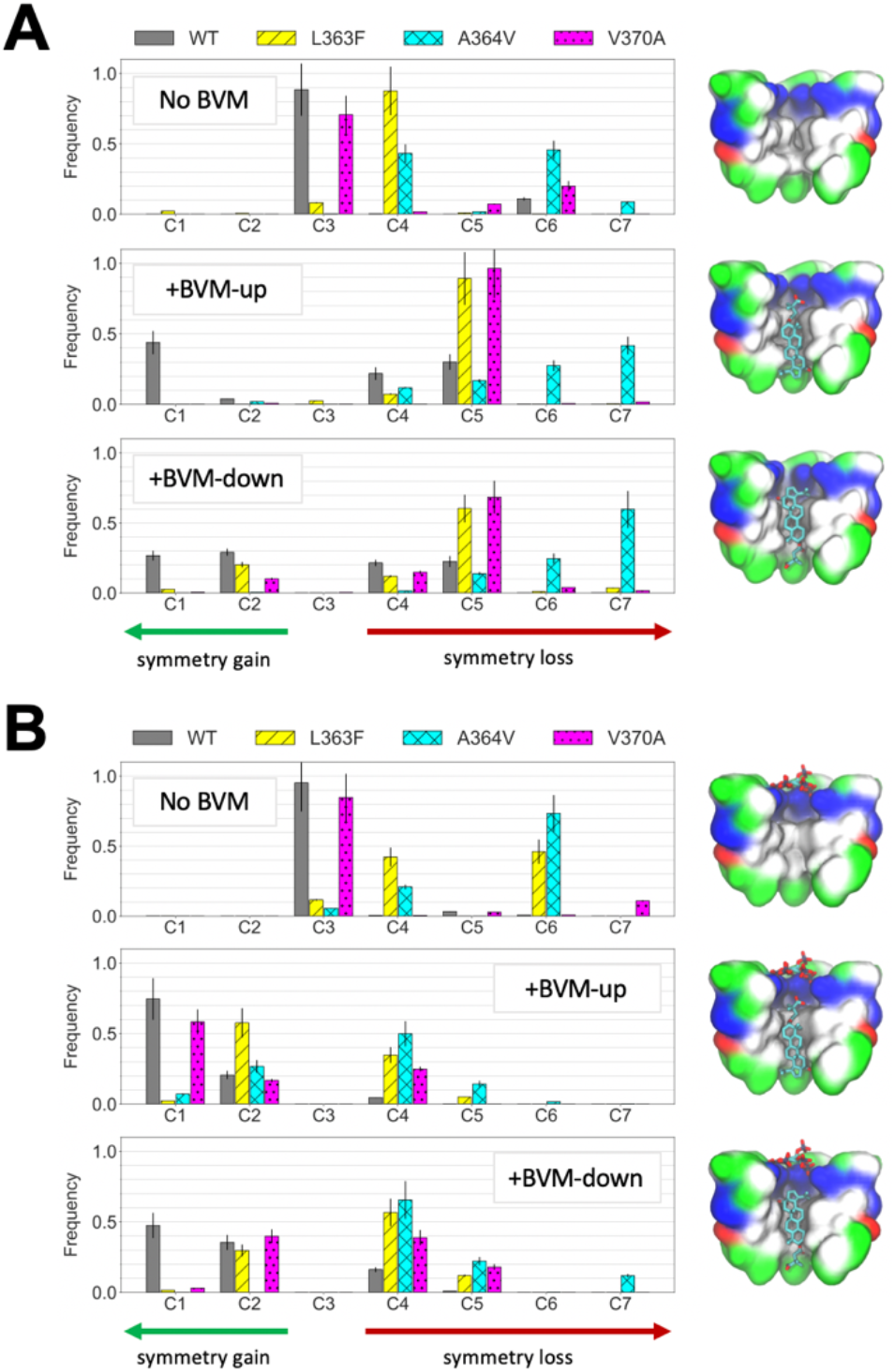
Comparison of cluster populations in WT and mutant (L363F, A364V, and V370A) viruses. The (A) absence and (B) presence of inositol hexakisphosphate (IP6) is compared in addition to the absence and presence of bevirimat (BVM) in two binding poses, BVM-up and BVM-down (n=17,274). The arrows at the bottom indicate the direction of symmetry gain (green) or loss (red) with respect to BVM-free WT virus. The schematics to the right show each system configuration as a cutaway, with half of the helical bundle as a colored surface (white is hydrophobic, green is polar, blue is positively charged, and red is negatively charged). BVM in the central pore is displayed in stick representation.

The three mutant CA_CTD_/SP1 hexamers exhibited varying levels of distortion to the 6HB in the absence of BVM. As seen in Figure 3A, taking increasing cluster indices as indicative of increasing loss of symmetry, we find that the preservation of 6HB structural integrity in the majority population varies approximately as WT > V370A > L363F > A364V. Note that all three mutant viruses express replication-competent phenotypes (*13, 14*), suggesting that the observed fluctuations in the 6HB structure do not impede viral assembly and maturation. However, the increased flexibility in the 6HB, most apparent in A364V, may facilitate protease access to the CA/SP1 scissile bond. Indeed, in the case of A364V, enhanced proteolytic cleavage of CA/SP1 compared to WT has been observed (*14, 15, 30*). By contrast, L363F and V370A had no discernible effects on proteolytic cleavage efficacy (*14, 30, 31*).

The addition of BVM has both mutant-specific and binding-pose specific effects on cluster populations. The BVM-up pose does not significantly increase the population of six-fold (cluster 1) or partially six-fold (clusters 2 to 4) symmetric clusters in the three mutant viruses. Instead, a broad distribution is observed with the majority population of L363F, A364V, and V370A 6HB states in low-symmetry clusters (clusters 5 to 7). These results suggest that the BVM-up pose is unable to recover the high-symmetry states that are stabilized by BVM in WT virus, and instead may have an antagonistic effect. However, the BVM-down pose shifts the cluster populations of L363F and V370A toward clusters 2 to 4, i.e., partially six-fold symmetric states. The implication would be that the BVM-down pose may partially recover the activity of BVM seen in WT virus. In the A364V case, however, the majority population remains in clusters 5 to 7, i.e., low-symmetry states. Both BVM binding poses appear to be ineffective against A364V.

Prior assays have shown that the fold change (FC) in half-maximal effective concentration (EC_50_), i.e., the ratio of EC_50_ in mutant viruses to that of WT virus, is 20-120 for V370A and >1000 for A364V (*32-35*). In separate CA/SP1 processing assays in the presence of BVM, the majority of CA/SP1 molecules were cleaved in A364V mutants, intermediate levels were cleaved in L363F and, to a lesser extent, V370A mutants, and small fractions were cleaved in WT virus (*14, 15, 30*). These assays imply that the experimental efficacy of BVM in blocking protease activity varies as WT > V370A > L363F > A364V. On the other hand, our cluster analysis implies that BVM-induced symmetry preservation varies as WT > L363F > V370A > A364V. Therefore, the ability of BVM to restrict the occurrence of low-symmetry 6HB configurations correlates with the efficacy of the drug, with the exception of L363F compared to V370A. We hypothesize that additional factors may influence BVM efficacy, and explore one such possibility next.

### Inositol hexakisphosphate (IP6) has cooperative effects with BVM on six-helix bundle stabilization

IP6 is an essential cofactor in HIV-1 replication and has been shown to bind to the aforementioned K359 ring in immature WT virus (*23*). Prior MD simulations indicate that IP6 has a stabilizing effect on the 6HB (*23*), similar to the observed effects of BVM. It is therefore of interest to investigate potential cooperative effects when both IP6 and BVM are present.

According to our cluster analysis, as seen in Figure 3B, the addition of IP6 to WT virus in the absence of BVM increases the population in majority cluster 3 by around 5%. However, cluster 1 remains unpopulated, indicating that IP6 itself does not induce the same degree of symmetry preservation as BVM (compare to Figure 3A). The addition of both IP6 and BVM, however, increases the population in cluster 1 to around 75% and 50% for BVM-up and BVM-down, respectively, an increase of around 35% and 25% compared to BVM by itself. The significant increase in the cluster 1 population suggests that the presence of IP6 and BVM within the central pore of the 6HB cooperatively promote six-fold symmetric states. It is also interesting that this effect is more pronounced in the BVM-up case. In this orientation, the dimethylsuccinyl moiety interacts with the K359 ring that also interacts with IP6. Rather than competing, both molecules appear to share the K359 binding sites, with the dimethylsuccinyl moiety of BVM and IP6 coordinating 1-2 and 4-5 lysine side chains, respectively (see Supplementary Figure 2).

Several mutant-specific and binding-pose specific effects on cluster populations also emerge when IP6 and BVM bind to the 6HB. In the absence of BVM, both L363F and A364V mutants populate lower symmetry states, primarily clusters 4 and 6, while the V370A mutant primarily populates cluster 2, the same as WT virus. Hence, IP6 has varied effects on mutant viruses (Figure 3B): IP6 increases 6HB symmetry preservation for V370A, while decreasing the same for L363F and A364V mutants. However, the resultant trend in 6HB symmetry preservation remains the same as IP6-free virus: WT > V370A > L363F > A364V.

All three mutants exhibit greater symmetry preservation in the 6HB when BVM is added to IP6-bound virus, although to varying degrees and more substantially in the BVM-up pose. The V370A mutant populates cluster 1 with around 60% for the IP6 and BVM-up case, while the L363F and A364V mutants populate cluster 2 with around 60% and 30%, respectively. For comparison, the WT virus populates cluster 1 with around 75% probability, indicating a loss of symmetry upon mutation, which is to be expected. Instead, the V370A mutant primarily populates cluster 2, the L363F mutant primarily populates cluster 3, and the A364V mutant primarily populates cluster 4. Taken together, these results show that the efficacy of symmetry preservation by IP6 and BVM-up varies as WT > V370A > L363F > A364V, which is consistent with prior FC-EC_50_ measurements and CA/SP1 processing assays (*14, 15, 30, 32-35*). Note that the former two mutants exhibit greater ordering compared to BVM-free WT virus while the latter exhibits greater disordering. These general trends are preserved in the BVM-down case, but the specific clusters that are populated by each mutant tend to be those with lower symmetry compared to the BVM-up case. Importantly, our analysis shows that IP6 cooperatively affects the activity of BVM and its role in experimental assays should be clarified in future maturation inhibitor studies.

### Escape mutants exhibit multiple mechanisms of failure

Having analyzed the effect of escape mutants on cluster populations with varying degrees of six-fold symmetry, we now investigate the origin of these observed differences. In particular, we quantify intermolecular contacts between BVM and hydrophobic residues along the interior cavity of the 6HB, as depicted in Figure 4A. We narrow our analysis to clusters 1, 2, 4, and 5 to provide a representative selection of 6HB configurational states upon BVM binding. In the following, we analyze conditions in which each of these clusters is highly populated, i.e., in the presence of BVM in the BVM-up orientation; IP6 is also included for clusters 1, 2, and 4.

**Figure 4.**
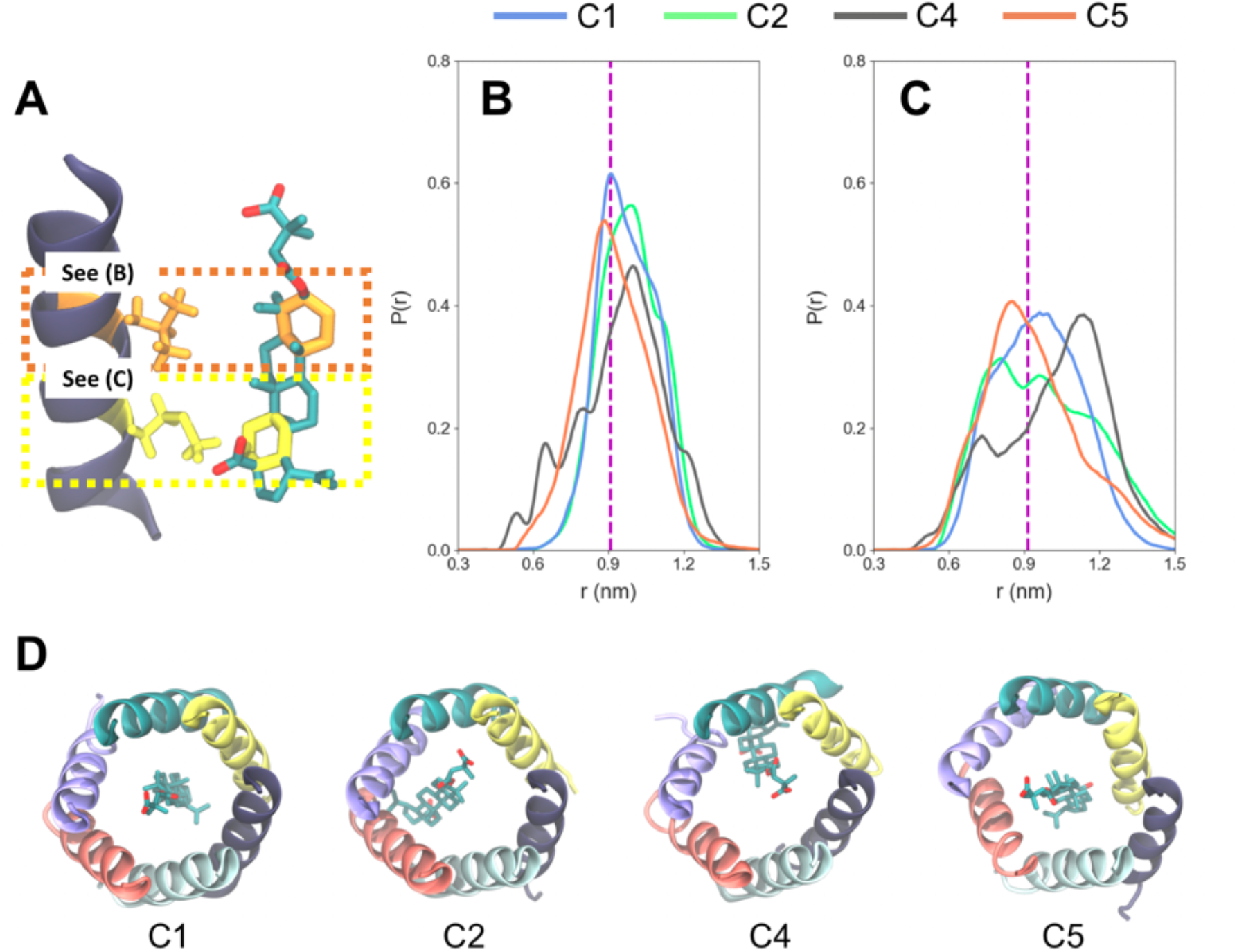
Proximity analysis between bevirimat (BVM) and the six-helix bundle (6HB). (A) Schematic showing the contacts that were measured between the upper ring of BVM and residue 363 within the 6HB (both orange) and the lower ring of BVM and residue 367 (both yellow). (B,C) Probability distributions (P(r)) of the radial distance (*r*) between the C atoms throughout the highlighted rings in the pentacyclic triterpenoid motif of BVM and C_α_ atoms of (B) L/F363 and (C) M367 within the listed cluster index (i.e., for clusters 1, 2, 4, and 5). The dashed line indicates the mean *r* between the respective moieties from the atomic model derived by MicroED (*18*). (D) Snapshots showing representative positions of BVM (stick representation) within the central pore of the 6HB (ribbon representation).

Using the cluster labels assigned to each configuration, we aggregated ensembles of each of the aforementioned clusters across WT and mutant virus trajectories for further analysis. Figure 4B-C show the computed probability distributions (P(*r*)) of the distance (*r*) between C atoms in the pentacyclic triterpenoid moiety of bevirimat and the Cα atoms of L363 (or F363 in the mutant case) and M367. Hereafter, we refer to these distances as *r*_363_ and *r*_367_, respectively; note that we use the Cα backbone as our reference point since the side chains will vary depending on the mutant.

The *r*_363_ and *r*_367_ distributions for cluster 1 (blue lines in Figure 4B-C), which represents six-fold symmetric 6HBs, are unimodal with peak positions that correlate with the mean *r* values from the experimental atomic model (*18*). The unimodality of the distribution indicates that the pentacyclic triterpenoid moiety is uniformly coordinated by each of the six helices, as seen in Figure 4D. Note that the broadening of the *r*_367_ distribution as compared to that of *r*_363_ suggests that while BVM uniformly interacts with L363 throughout the six helices, BVM may preferentially interact with a few M367 residues throughout the six helices, which is likely due to favorable hydrophobic interactions.

For cluster 2 (green lines in Figure 4B-C), the *r*_367_ distribution is multimodal with the largest peak shifted to smaller *r* compared to the experimental *r*_367_ value. The *r*_363_ distribution, on the other hand, remains largely unimodal. In this cluster, BVM appears to closely interact with a subset of the helices around M367 while remaining uniformly coordinated by L/F363; in other words, BVM tilts such that the bottom of the pentacyclic moiety establishes closer contact with the 6HB (see Figure 4D). For cluster 4 (grey lines in Figure 4B-C), the *r*_363_ and *r*_367_ distributions exhibit multimodality with peaks appearing below the experimental *r*_363_ and *r*_367_ values. Here, the entire pentacyclic moiety of BVM appears to establish close contacts with a subset of helices around L/F363 and M367. Recall that clusters 2 and 4 both represent 6HBs with partial six-fold symmetry, suggesting that BVM fails to maintain six-fold symmetry due to preferential association with a subset of helices that contain escape mutations. This behavior is notable for V370A and L363F mutants.

The *r*_363_ and *r*_367_ distributions for cluster 5 (red lines in Figure 4B-C) are unimodal, yet are broadened with respect to the cluster 1 distributions. The broadening of the *r*_363_ distribution indicates that the L/F363 coordination around the pentacyclic triterpenoid moiety has increased flexibility. Both the broadening and extended tail of the *r*_367_ distribution is significant as it suggests that the 6HB near M367 exhibits increased flexibility commensurate with weak (or non-specific) association between BVM and M367. Note that similar tail behavior is seen in the cluster 2 and 3 distributions.

### Loss of six-fold symmetry in the six-helix bundle increases uncoiling propensity

To further quantify the effect of BVM, we compute free energy surfaces (FES) that describe the helix-to-coil transition in the 6HB. Here, we use well-tempered metadynamics (*36*), a technique that dynamically adds external bias to the system to promote rare event sampling, from which the FES is recovered through reweighting (see Methods). We project the FES along two coordinates, Ψ_6_^2D^ and the so-called alpha-beta similarity (*AB*_*sim*_) (*37*), which assesses the collective deviation of *ϕ* and *Ψ* torsional angles from that of α-helices:

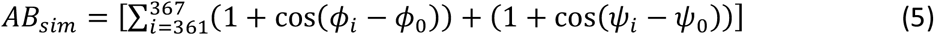

where *ϕ*_*i*_ (*Ψ*_*i*_) is the N-Cα (Cα-C) backbone torsional angle of residue *i* and *ϕ*_*0*_ (*Ψ*_*0*_) is the reference angle (−60° for both) for α-helices; we focus on residues 361 to 367 as the scissile bond is located between residue 363 and 364. When the torsional angles throughout the measured protein backbone are equivalent to α-helices, *AB*_*sim*_ is expected to be 14 (twice the number of residues) and linearly diminishes toward 0 with increasing deviation, i.e., uncoiling.

We compare the FES computed for WT virus with IP6 (Figure 5A-B) to that of WT virus with IP6 and BVM (Figure 5C-D) and L363F mutant virus with IP6 and BVM (Figure 5E-F). In WT virus with IP6, two primary free energy basins are observed. The first is the folded α-helical state around *AB*_*sim*_ = 14 and Ψ_6_^2D^ = 0.60, which transitions to an uncoiled state that broadly spans a range of 4 < *AB*_*sim*_ < 12 with a free energy barrier height of 2.5 kcal/mol (see Figure 5B). Note that the uncoiled state also spans a broad range of 0.35 < Ψ_6_^2D^ < 0.70 with a local free energy minimum around Ψ_6_^2D^ = 0.50. Hence, helical unfolding is commensurate with a reduction in 6HB symmetry.

**Figure 5.**
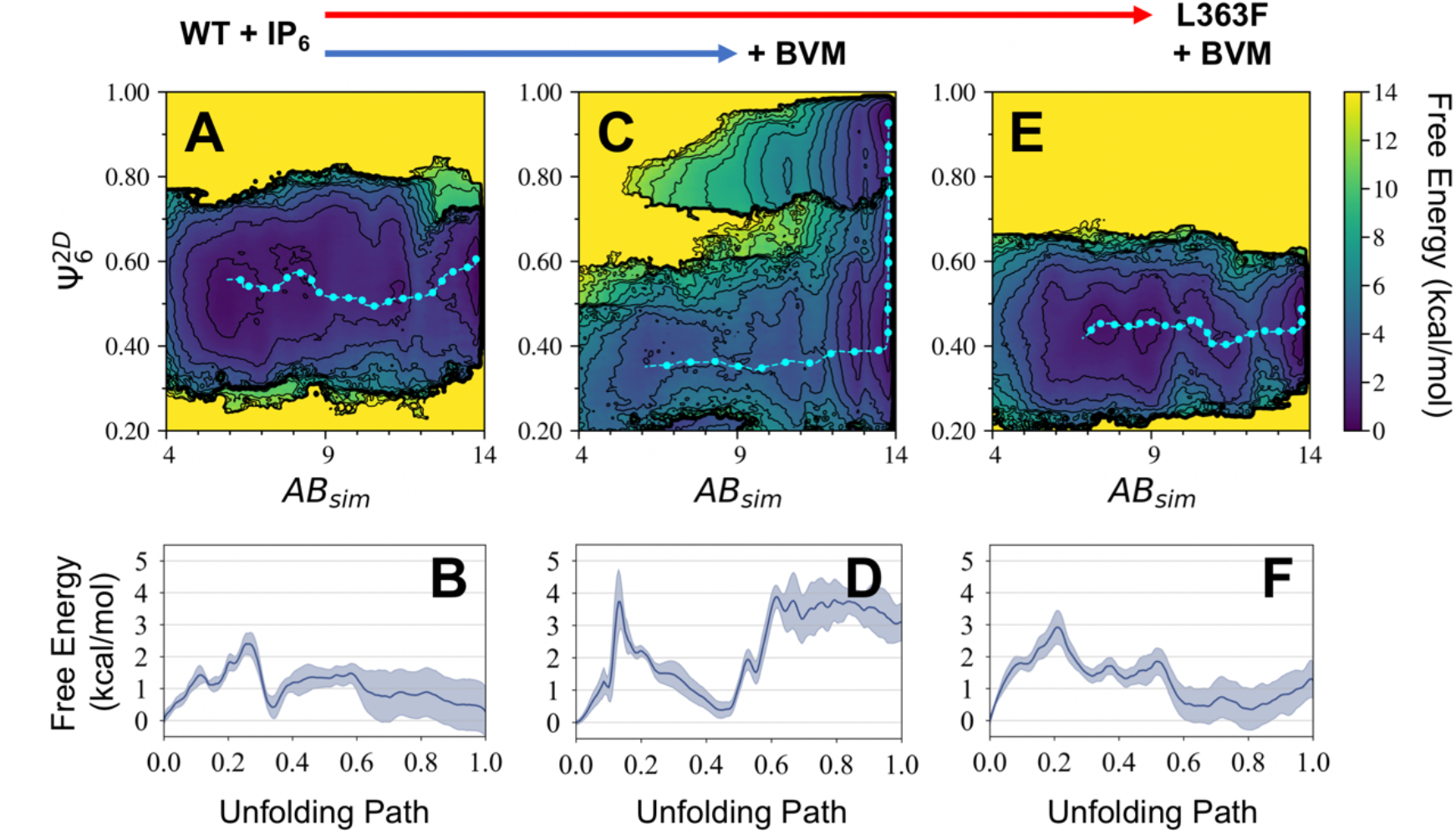
Free energy surfaces for helix-to-coil transitions. Comparison between CA_CTD_ /SP1 for (A-B) wild-type (WT) with IP6, (C-D) WT with IP6 and BVM, and (E-F) L363F mutant with IP6 and BVM. (A,C,E) Free energies are projected onto two coordinates: alpha-beta similarity (*AB*_*sim*_), which describes the extent the region around the proteolytic cleavage site (i.e. L363/A364) is folded into an α-helix, and the six-fold Mermin order parameter (Ψ_6_^2D^), which describes the degree of six-fold symmetry throughout the six-helix bundle. (B,D,F) 1D minimum free energy paths along the dotted cyan lines in (A,C,E) that describe helix uncoiling (n=3 per panel); the start of the unfolding path corresponds to the start of the dotted cyan line when *AB*_*sim*_ ≅ 14.

As seen in Figure 5C, the addition of BVM to WT virus with IP6 introduces another basin representing the folded state (*AB*_*sim*_ = 14) in a high-symmetry 6HB (Ψ_6_^2D^ = 0.9). For uncoiling, two structural transitions occur in sequence. The 6HB first distorts into a lower symmetry state (Ψ_6_^2D^ = 0.40) with a barrier height of 3.5 kcal/mol, followed by uncoiling with a second barrier height of 3.0 kcal/mol (see Figure 5D). The distortion of the 6HB represents a significant barrier against uncoiling introduced by BVM. In contrast, this high-symmetry state is abolished in the L363F mutant with IP6 and BVM, as seen in Figure 5E. In this case, note that the 6HB tends to adopt low-symmetry configurations similar to that of WT virus. The barrier height for uncoiling, however, is 3.0 kcal/mol (see Figure 5F), an increase of 0.5 kcal/mol compared to that of WT virus, suggesting that the presence of BVM slightly decreases the propensity for uncoiling despite the symmetry loss throughout the 6HB. Nonetheless, it is clear from our analysis that BVM primarily prevents helical uncoiling by stabilizing high-symmetry 6HB states.

## Discussion

Taken together, our analysis suggests that BVM coordinates the 6HB in WT virus, thereby abrogating helical uncoiling, inhibiting protease access, and preventing maturation. To effectively coordinate the 6HB uniformly, BVM stochastically rotates within the bundle, hopping from protomer to protomer. Our simulations also suggest that both BVM-up and BVM-down are viable binding poses, although we predict that BVM-up is the more effective inhibitor. In mutant viruses, however, we observe two primary structural states that are related to their escape mechanisms, and provide the following interpretation that is depicted using helical wheel diagrams in Figure 6.

**Figure 6.**
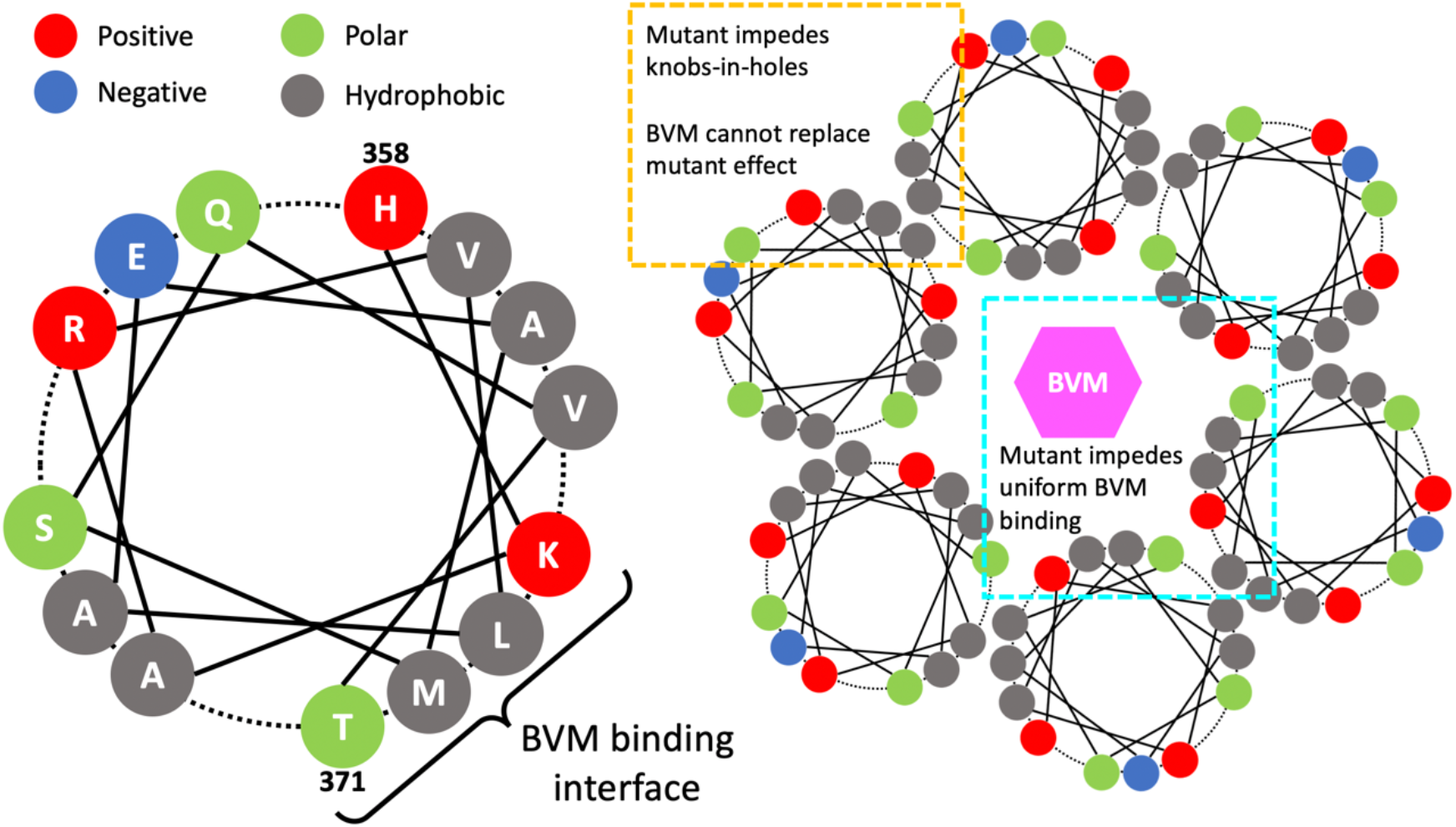
Schematic of bevirimat (BVM) and escape mechanisms using helical wheel diagrams. (Left) depiction of the CA_CTD_/SP1 junction helix (H358 to T371) and (right) of the six-helix bundle (6HB); the color of each residue label indicates its type: positively charged, negatively charged, polar, or hydrophobic. In the 6HB projection, the grey circles in the orange box establish hydrophobic knob-in-hole (KIH) interactions that stabilize 6HB packing. The A364V and V370A mutants target these interactions and negatively affect KIH packing, which BVM does not compensate. The grey circles in the cyan box establish hydrophobic interactions with BVM. The L363F mutant targets these interactions and negatively affects uniform BVM coordination by the 6HB.

First, escape mutations can disrupt the hydrophobic binding interface along the interior of the 6HB (cyan box in Figure 6), i.e., L363 and M367. According to the cluster 1 distributions in Figure 4B-C, BVM interacts more strongly with L363 compared to M367. Given the larger hydrophobicity of phenylalanine compared to leucine (*38, 39*), the L363F mutant likely increases the microscopic binding affinity of BVM. Yet, due to the flexibility of phenylalanine, we find that BVM preferentially coordinates to a subset of phenyl groups; in our simulations, we see these phenyl groups flatten, thereby establishing contacts with the pentacyclic moiety, while the remaining phenyl groups extend into the vacated space within the central pore and the 6HB distorts accordingly. To improve BVM coordination, these key residues likely need to be mutated to hydrophobic residues with stiff side chains, such as branched amino acids. For instance, M367I has been shown to have high BVM efficacy (*40*).

Second, escape mutations can disrupt the hydrophobic knob-in-hole (KIH) interfaces (*16*) that are essential to the quaternary structure of the 6HB (orange box in Figure 6). Here, the “knobs” consist of methyl side chains from A360 and A364 that coordinate within “holes” that are formed between side chains from the adjacent helix (the next counterclockwise helix in the 6HB in Figure 6). These side chains are composed of V362, L363, and V366 for A360 and A366, M367, and V370 for A364. The asymmetry between the knob and hole side chains is likely important as the smaller methyl knobs can be coordinated by the bulkier side chains that form the hole. Hence, increasing the knob size with A364V mutants or decreasing the hole size with V370A mutants lead to loss of KIH packing efficacy, resulting in loss of 6HB symmetry. In these cases, we find that BVM interactions with the interior hydrophobic interface of the 6HB (cooperatively enhanced when IP6 is present) is insufficient to retain 6HB symmetry.

Although we did not simulate it, mutants may instead stabilize the 6HB such that BVM has negligible perceived efficacy. In this case, mutant virus in the absence of BVM can sufficiently symmetrize the 6HB to prevent protease access. On the basis of the aforementioned KIH interactions, one way to do so may be to mutate hole-forming residues with branched (i.e., bulkier and stiffer) and more hydrophobic side chains, e.g., valine, leucine, and isoleucine. Indeed, L363I, A366V, and M367I mutants have been shown to decrease viral infectivity and CA/SP1 processing, while BVM itself had indiscernible effects on L363I mutant virus (*14, 40*). The T371I mutant is also known to prevent maturation and phenocopies BVM (*19, 41*), while BVM likely induces no additional symmetry preservation in the 6HB.

Our findings highlight difficult constraints that must be overcome by next-generation maturation inhibitors, which includes precise coordination to the 6HB in both WT and polymorphic viruses. The dimethylsuccinyl moiety has been recognized as a pharmacophore (*32, 42*) for BVM, and our analysis shows that its coordination to K359 has potent 6HB symmetry preservation effects. Interestingly, this effect is cooperatively enhanced by the presence of IP6, which suggests modification of the dimethylsuccinyl moiety to improve overall K359 coordination may be valuable. The recognition that the QVT motif is a major source of viral polymorphism has motivated numerous modifications near the C-28 carboxylic acid moiety (*30, 32, 33, 35, 43, 44*), which have demonstrated that it is possible to overcome loss of susceptibility due to polymorphism. On the basis of our results, we suggest that modification or replacement of the pentacyclic triterpenoid moiety is a potential third avenue. As this moiety establishes key contacts with the hydrophobic interior of the 6HB, it may be possible to emulate the effect of mutants that simultaneously stabilize KIH and inhibitor interactions. For instance, a triterpenoid with larger hydrophobic (e.g. propyl) groups near the terminal cyclic rings may aid in M367 interactions and facilitate downstream KIH interactions. Finally, this work suggests the need to evaluate inhibitor activity from the perspective of structural ensembles, and to that end, we anticipate that molecular simulations will be a valuable tool for fundamental inhibitor studies and design.

## Conclusions

Using extensive MD simulations and free energy sampling, we have provided a comprehensive comparison of molecular structure-activity relationships across WT HIV-1 and mutant (L363F, A364V, and V370A) CA_CTD_ /SP1 viruses in the presence and absence of BVM, a maturation inhibitor. Our analysis reveals an ensemble of 6HB structures with varying degrees of six-fold symmetry. The most important conclusion is that the inclusion of BVM in WT virus rigidifies the 6HB thereby preserving six-fold symmetry, which is cooperatively enhanced by IP6, the assembly cofactor. The escape mutants reduce six-fold symmetry throughout the 6HB, which is partially recovered by BVM in particular cases. Our free energy calculations demonstrate that loss of six-fold symmetry facilitates helical uncoiling, which precedes proteolytic cleavage (*19, 22*). Consequently, the efficacy of BVM is due to additional resistance to symmetry loss. Two primary escape mechanisms were identified: (i) failure to mitigate lost knob-in-hole 6HB interactions and (ii) failure to uniformly coordinate the 6HB within the central pore. Our analysis demonstrates the utility of MD simulations to provide insight into the molecular basis for the stabilization of the CA_CTD_ /SP1 six-helix bundle conferred by BVM, which provides new directions for the design of future HIV maturation inhibitors.

## Methods

All MD simulations were prepared using hexameric CA_CTD_/SP1 domains of an atomic model (PDB: 5L93) (*17*), which was modified accordingly with the point mutations L363F, A364V and V370A. Systems prepared with IP6 and BVM were initialized with Monte Carlo insertion of the molecule in the central pore of the hexamer near K359 and M367, respectively. Each system was solvated by water in a dodecahedral domain with at least 1.6 nm of space between the edges of the protein and the domain. Water molecules were then randomly replaced with 0.15 M of NaCl. The total size of each system was ∼108,000 atoms with initial box lengths of 11.6×11.6×8.2 nm.

All simulations used the CHARMM36m force field (*45*) and were performed using GROMACS 2016 (*46*). Minimization was performed using steepest descent until the maximum force was reduced to 1000 kJ/mol/nm. Then, equilibration was performed in several phases. First, 10 ns were integrated in the constant *NVT* ensemble using the stochastic velocity rescaling thermostat (*47*) with a damping time of 0.2 ps and a timestep of 1 fs. During this phase, the Cα backbone of the protein was harmonically restrained with a force constant of 1000 kJ/mol/nm^2^. An additional 40 ns were integrated in the constant *NPT* ensemble using the stochastic velocity rescaling thermostat (*47*) (2 ps damping time) and the Parrinello-Rahman barostat (*48*) (10 ps damping time) and a timestep of 2 fs. Finally, an additional 750 ns were integrated in the constant *NVT* ensemble using the Nose-Hoover chain thermostat (*49*) (2 ps damping time) and a timestep of 2 fs. During the latter three phases, the Cα backbone of helix 9 throughout the hexamer was harmonically restrained with a force constant of 500 kJ/mol/nm^2^. Throughout this procedure, H-containing bonds were constrained using the LINCS algorithm (*50*). Statistics were gathered every 200 ps, and the final 600 ns of each trajectory was used for analysis, which were performed using a combination of VMD Tcl scripts (*51*) and Python scripts using MSMBuilder (*52*) and MDTraj (*53*).

Free-energy calculations were performed using PLUMED 2.4 (*54*) and GROMACS 2016 (*46*). Final configurations from the MD simulations for each of the three reported systems were used as initial configurations. For each system, three independent replicas were prepared using random velocity distributions and equilibrated in the constant *NVT* ensemble (same thermostat and timestep as above) for 200 ns; the same restraints on helix 9 and H-containing bonds were kept throughout these simulations. Well-tempered metadynamics (*36*) was subsequently performed for 4.6 µs by biasing the *AB*_*sim*_ collective variable for a single helix; Gaussian hills with 0.5 kJ/mol height and 0.05 width were deposited every 1 ps with a bias factor of 15 *k*_*B*_*T*. Data was collected every 1 ps and the Tiwary-Parrinello algorithm (*55*) was used to reweight FESs onto a 2D projection along *AB*_*sim*_ and Ψ_6_^2D^. Here, Ψ_6_^2D^ was computed using the non-biased helices, i.e., only four (out of six) angles were computed and averaged, as the biased helix twists out of the 6HB when unfolding. All reported FESs were averaged across replicas after aligning free energy values at the observed global minimum.

## Supplementary Information

Two additional figures, one showing a comparison of α-helix similarity of the C_α_ backbone (*α*_*sim*_) throughout the six-helix bundle (6HB) for each cluster, and another showing a representative schematic of BVM and IP6 coordinating to residue K359 within the 6HB, can be found in the Supplementary Information.

## Acknowledgments

This research was supported in part by the National Institute of Allergy and Infectious Diseases (NIAID) of the National Institutes of Health under grant R01 AI154092 (for M.Y. and G.A.V.). A.J.P. acknowledges support from the NIAID under grant F32 AI150477. The authors are grateful to Dr. Aleksander Durumeric for useful discussions and comments on the manuscript. All computational work was performed using Department of Defense (DoD) resources as part of the High-Performance Computing Modernization Program (HPCMP).

## Author Contributions

A.J.P., M.D.P., M.Y., and G.A.V. conceived the study and wrote the manuscript. A.J.P. performed simulations. M.D.P provided experimental data. Data analysis was performed by A.J.P., M.D.P., M.Y., and G.A.V.

## Data Availability

The data that support the findings of this study are available from the corresponding author upon reasonable request.

