## Supplemental File for "Preservation of HIV-1 Gag helical bundle symmetry by bevirimat is central to maturation inhibition"

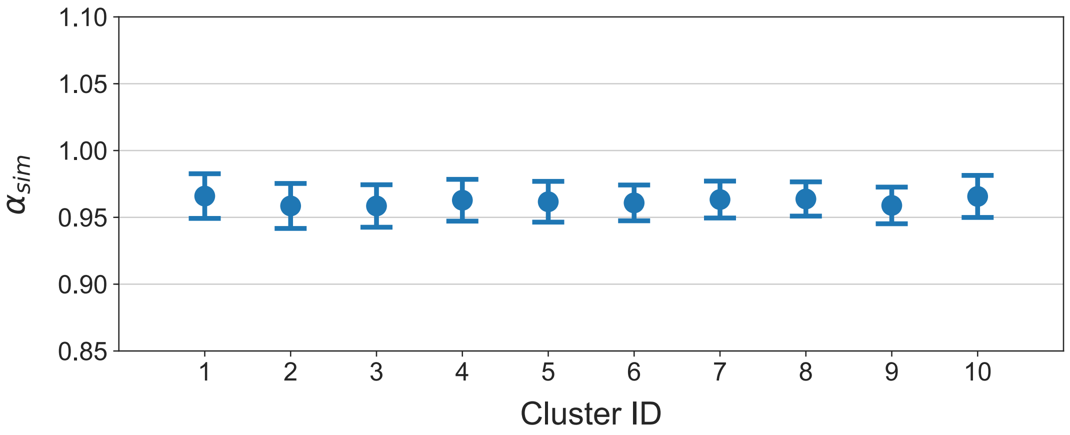


**Supplementary Figure 1**. Comparison of α-helix similarity of the $C_\alpha$ backbone ($\alpha_{sim}$) throughout the six-helical bundle (6HB) for each cluster (see Main Text). The α-helical character in the 6HB is largely preserved and consistent across clusters.

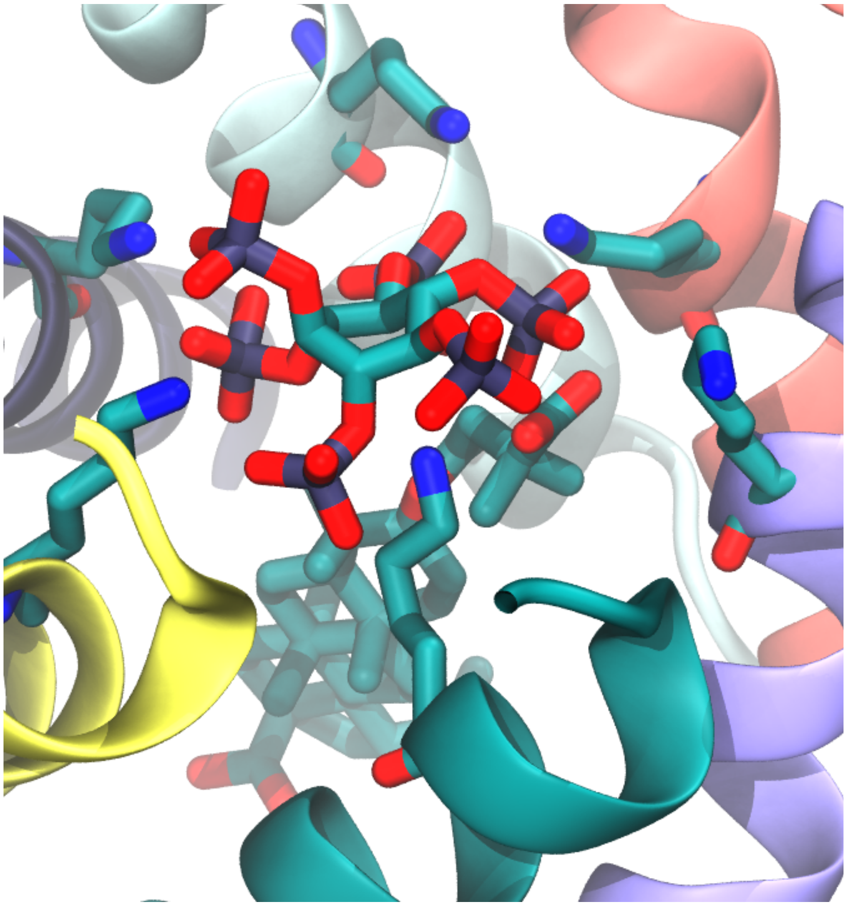

**Supplementary Figure 2**. Representative schematic of BVM and IP6 coordinating to K359 within the six-helical bundle. To accommodate coordination between the BVM carboxylic moiety and a lysine sidechain, IP6 tilts yet remains coordinated up to five lysine sidechains.
